# A high-throughput method for unbiased quantitation and categorisation of nuclear morphology

**DOI:** 10.1101/312470

**Authors:** Benjamin Matthew Skinner, Claudia Cattoni Rathje, Joanne Bacon, Emma Elizabeth Philippa Johnson, Erica Lee Larson, Emily Emiko Konishi Kopania, Jeffrey Martin Good, Gullalaii Yousafzai, Nabeel Ahmed Affara, Peter James Ivor Ellis

## Abstract

The physical arrangement of chromatin in the nucleus is cell type and species specific. This is particularly evident in sperm, in which most of the cytoplasm has been lost; the shape of the nucleus reflects the shape of the cell. Mice have distinctive falciform (‘hook shaped’) sperm heads and nuclei. Quantification of the differences in shape variation between mouse species and lines often relies on manual measurement and classification that leads to subjective results, making comparisons within and between samples difficult.

We have developed an analysis program for assessing the morphology of asymmetric nuclei, and characterised the sperm of mice from a range of inbred, outbred and wild-derived mouse lines. We find that laboratory lines have elevated sperm shape variability both within and between samples in comparison to wild-derived inbred lines, and that sperm shape in the F1 offspring of CBA and C57Bl6J lines is subtly affected by the direction of the cross.

Hierarchical clustering can distinguish distinct sperm shapes with greater efficiency and reproducibility than even experienced manual assessors. We quantified the range of morphological defects in the inbred BALB/c line, demonstrating we can identify different morphological subgroups. This approach has applications for studies of sperm development, infertility and toxicology.

## Introduction

Nuclei are complex, dynamic structures within a cell. For many cell types, the nucleus is generally spherical, but for other cell types the nucleus adopts a distinctive shape [1]. One of the most profound changes to nuclear shape occurs during spermatogenesis: mammalian sperm tend to have a spatulate, or ‘paddle’ shape, meaning the nucleus both condenses and reshapes. The chromatin becomes wound ~4-6 times more tightly than in metaphase, mediated via replacement of histones with smaller protamines [2], and various cytoskeletal elements coordinating to shape the nucleus [3].

In rodents, this process is even more elaborate: most rodents, including mice, have a falciform ‘hook-shaped’ sperm, with varying degrees of hook length and body shape between species (e.g. [4]). The mouse sperm head shape develops through a series of interacting mechanical forces, reshaping the nucleus via the cytoskeleton and nucleoskeleton. The sperm head is divided into developmental ‘modules’, each of which is shaped by particular cytoskeletal components [5]. When these processes go awry, distinct morphological abnormalities can result (e.g. [6]), linking phenotype with the underlying genetic alterations. The reshaping of the nucleus is itself a distinct process from the chromatin condensation and repackaging [3]. Reshaping precedes transition and protamine replacement, and chromatin condensation then follows.

Analytical methods for categorising and quantifying sperm head shape variation have developed markedly over the years, and the advent of computational processing of images has dramatically increased the quality of data we can capture, and the sophistication of the analyses. To date, morphometric approaches in sperm have fallen into three main groups; the measurement of basic parameters such as lengths, widths, and areas of objects, the use of elliptic fourier analysis to investigate differences in the two dimensional outline of the object, and the use of Procrustes analyses to examine differences in fixed landmarks within the sperm head. Each of these approaches has advantages and disadvantages.

Basic measures such as area and length were the first statistics recorded describing sperm morphology (e.g. [7–9]. These still remain useful, especially in situations such as CASA analysis for fertility screening, in which an assessment of semen quality must be made rapidly across many different cells [10]. However, the parameters measured by these analyses are dominated by the size of the object, not the shape, and can make it difficult to consistently assess the number of normal sperm across populations [11].

In contrast, elliptic Fourier descriptors [12] allow an arbitrary closed two dimensional shape to be decomposed into harmonic amplitudes describing the curvature of the object perimeter, thus allowing subtle variations in shape to be discovered [13]. This approach has proved powerful for demonstrating differences between species, lines within a species, and different treatments (e.g. [14–16]). However, the approach has the drawback that the shape parameters and underlying mathematics are difficult for biologists to understand and relate back to the biological structure that is affected [17]. Moreover, since Fourier analyses rely on smooth harmonic deformations of an underlying elliptical outline, sharp points - such as found at the tip of a mouse sperm - tend to be poorly fitted [18].

The third major method, Procrustes-based geometric morphometric analysis, uses landmarks and semilandmarks within the object to align individual samples to consistent size, position and orientation (e.g. [4]). Principal component analysis (PCA) can then be used to identify the major varying landmarks distinguishing samples [5]. This approach has the advantage of tightly relating the variation to physical structures within the object: however, since objects are aligned by a least-squares method rotating about the centroid, objects are susceptible to smearing of landmarks in highly variable regions, and can require time-consuming manual placement of landmarks.

In terms of the biological field of application, sperm shape analysis has proven useful in three main interrelated areas: infertility, speciation, and toxicology. In infertility, while abnormal sperm morphology is extremely common in infertile knockout lines, the role played by specific types and extents of shape defect remains to be elucidated, as does the extent to which teratozoospermia can be used as an indicator of other sperm defects (e.g. DNA damage or defective motility [19]). Deregulation of reproductive processes is a major contributor to speciation through the induction of hybrid male sterility [20]. In particular, sperm shape abnormalities are a feature of house mouse hybrid sterility, with a range of mapped quantitative trait loci known, particularly on the sex chromosomes but also on autosomes [21–24].

Sperm shape is used as an assessment of genotoxicity and/or reproductive toxicity of compounds (e.g. [25,26]. These studies often carry out a manual classification of sperm into various categories of morphological abnormality, based on previously described sperm shapes. The manual element thus makes this application both time consuming, and prone to operator bias. A further problem is that the classes of abnormality described are often arbitrarily chosen, and vary between studies. Use of a scoring chart, based on the morphological abnormalities typical for one experimental system, may therefore compromise the ability to quantitate abnormalities in a different system. It would be far more useful to have an automated and reproducible method that is able to discover categories of morphological abnormality within a sperm population, without prior training.

To address these needs for unbiased measurement, analysis and categorisation of nuclear morphologies, we have developed a new image analysis programme that generates quantitative information on the underlying regions of the nucleus that differ within and between samples, independent of nuclear size.

We have validated the software on different mouse lines, and can quickly analyse hundreds of images. Here, we demonstrate the use of this software to compare a range of different inbred, outbred and wild-derived lines (revealing the effects of inbreeding depression and potentially hybrid dysgenesis), to unravel the morphological variation in a single sample (revealing different classes of abnormality in an inbred line), and to trace genetic influences on sperm morphology in a reciprocal F1 cross between CB57Bl6 and CBA lines (revealing contrasting effects of the parental genomes on sperm size and shape).

## Methods

### Mouse lines

All animal procedures were in accordance with the United Kingdom Animal Scientific Procedures Act 1986 and the University of Montana Institute for Animal Care and Use Committee (protocol 002-13) and were subject to local ethical review. Animals were sourced as indicated in Table 1; either from an approved supplier (Charles River Laboratories, Manston, UK), bred at Cambridge University Central Biomedical Services (Home Office licenses 80/2451 and 70/8925 held by PE), or bred at the University of Montana. Breeding colonies at the University of Montana were established from mice purchased from Jackson Laboratories (Bar Harbor, ME) or were acquired from Francois Bonhomme (University of Montpellier). Animals were housed singly or in small groups, sacrificed via CO2 followed by cervical dislocation (UM) or only cervical dislocation and tissues collected *post mortem* for analysis.

**Table 1:**
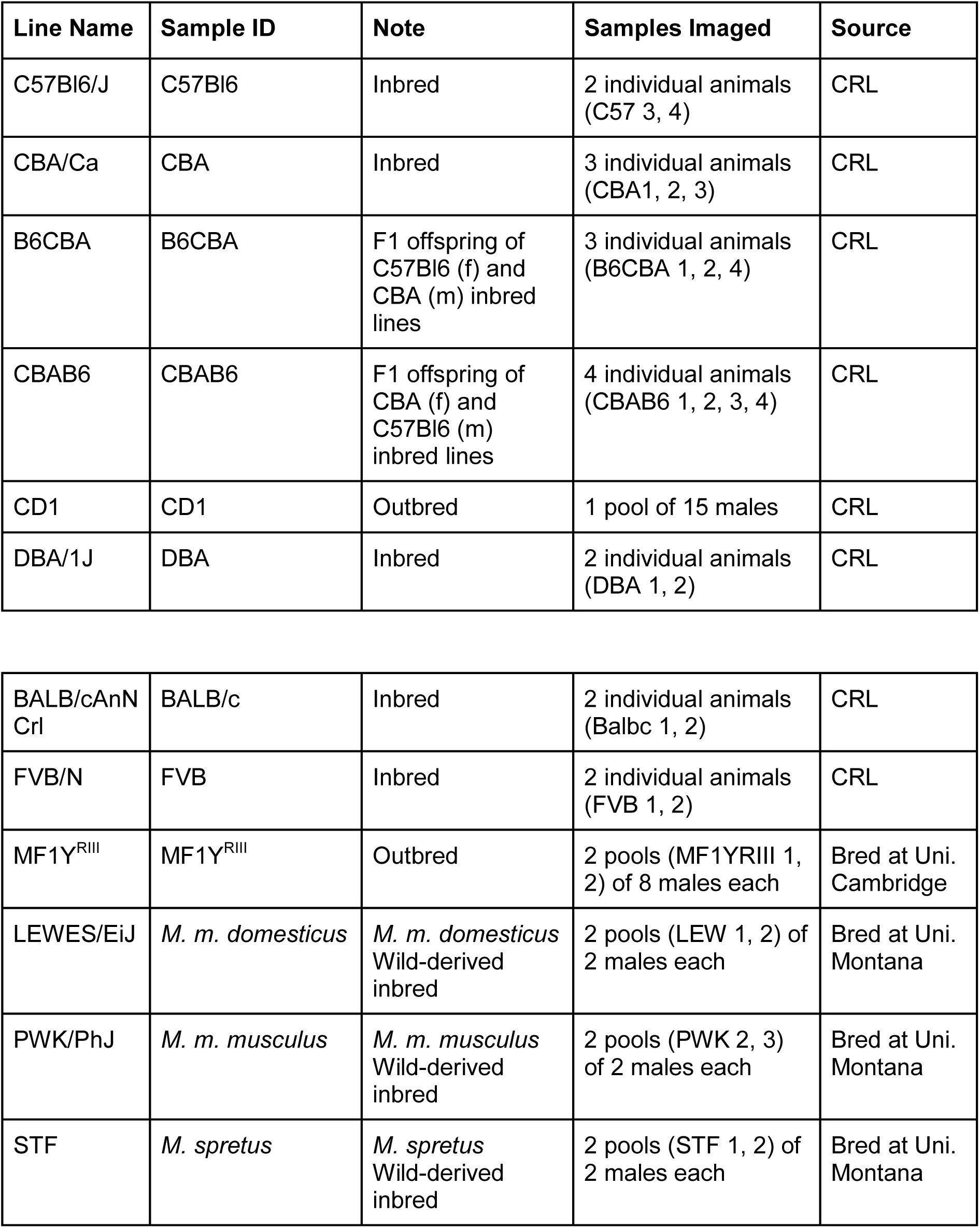
Mouse lines analysed for this study. CRL; Charles River Laboratories, Manston, UK.

### Sperm collection and fixation

The vasa deferentia and caudae epididymes were dissected from each animal, and the contents squeezed out into 1ml PBS (scaled up accordingly if multiple animals were pooled). The sperm were transferred to a microfuge tube, and tissue clumps were allowed to settle. Sperm were transferred to a new tube and pelleted at 500g for 5mins. The supernatant was removed, and the sperm fixed dropwise with either 3:1 methanol-acetic acid or 2% paraformaldehyde (PFA) in PBS. Sperm were again pelleted at 500g for 5mins, and washed in fixative twice more. Samples were stored at −20°C (methanol-acetic acid) or 4°C (PFA).

### Imaging

Samples were diluted in fixative as required to obtain an evenly-spread preparation, and 8μl of sample dropped onto a slide and allowed to air dry. Slides were counterstained with 16μl VectorShield with DAPI (Vector Labs) under a 22x50mm cover slip and imaged at 100x on an Olympus BX-61 epifluorescence microscope equipped with a Hamamatsu Orca-ER C4742-80 cooled CCD camera and appropriate filters. Images were captured using Smart-Capture 3 (Digital Scientific UK). To validate the reproducibility of the software, sample images were also gathered on three other microscopes: (1) an Olympus BX61 with a Hamamatsu C10600 orca r^2^ camera, (2) an Olympus BX61 with a Hamamatsu Orca-03G camera, and (3) a Nikon Microphot-SA epifluorescence microscope with a Photometrics Metachrome II CH250 cooled CCD camera.

### Nucleus detection and morphological analysis

Image analysis was performed using a custom program designed as a plugin for the freely available image analysis program ImageJ [27]. The plugin, Nuclear Morphology Analysis. The core software was developed using Java 8, with the user interface written using Swing. The software is available at http://bitbucket.org/bmskinner/nuclearmorphology/wiki/Home/ together with full installation instructions, an online wiki user manual, and example images for testing. The analyses described here were conducted using software version 1.13.6. The program allows for (a) detection of objects within fluorescence images and (b) morphological analysis of objects as sperm nuclei using a species-specific set of rules for identifying biologically relevant structures. The detection strategy is outlined in the supplementary methods.

Once nuclei were acquired from a set of images, they were consistently oriented and aligned. We used a modification of the Zahn-Roskies (ZR) transform [28] to ‘unroll’ the outline of each sperm nucleus by measuring the interior angle of the sperm at each point around the nuclear perimeter to generate a linear trace we refer to as the angle profile (Figure 1); further details and validation of the robustness of the method are given in the supplementary data and Supplementary figures 1-11.

**Figure 1:**
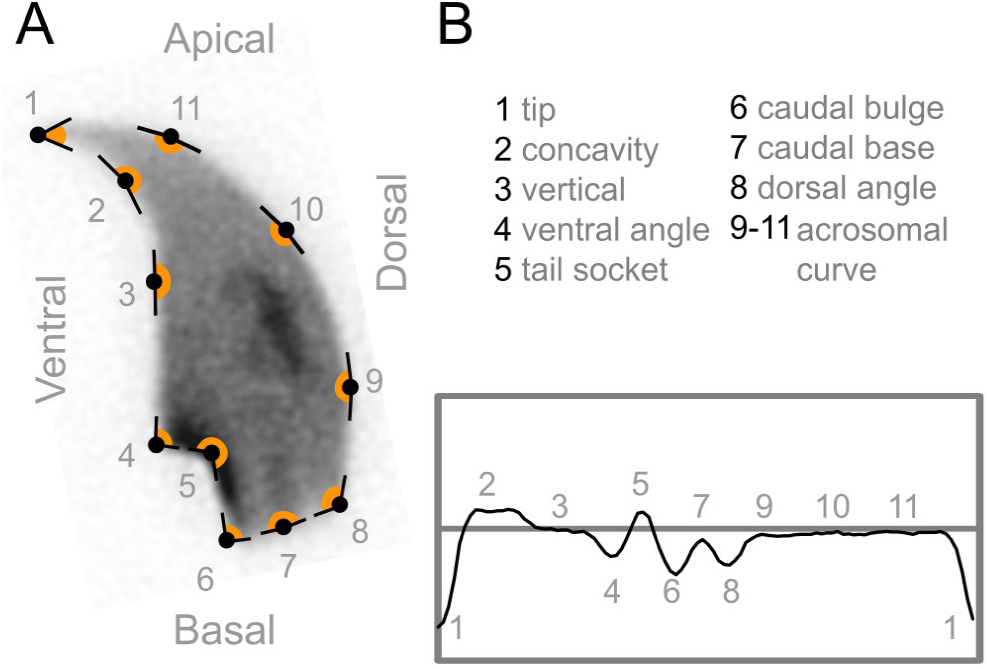
Shapes are detected by measuring the internal angles around the periphery of the nucleus. Angles from key features in an example nucleus (A) are plotted in (B). The actual profile for the entire perimeter is shown in (C).

### Statistical analysis and clustering

Following segmentation, standard nuclear parameters were measured: area, perimeter and aspect ratio, the width of the nuclear body versus the length of the hook as described in other papers (e.g. [9], and the lengths of each perimeter segment. Data was exported for further processing in R. Differences between datasets were tested using a pairwise Wilcoxon rank sum test, with Bonferroni multiple testing correction. In order to quantify the variability of the nuclear shapes, we developed a new per-nucleus measure defined as the root-mean-square difference between the per-nucleus angle profile and the median angle profile for the dataset, averaged across the length of the angle profile. The coefficient of variability (standard deviation / mean) was also calculated for each of the other measured parameters.

The ‘average shape’ of the nuclei was calculated by averaging the x and y coordinates at consistent semilandmarks taken as fractions of the perimeter across all nuclei, vertically aligned and with their centres of mass at (0,0). This yielded a ‘consensus nucleus’ visualising the overall shape of the population. Clustering was implemented via the WEKA data mining software library [29].

## Results

### Detection and quantification of sperm shape in C57Bl6 and CBA mice

The difference between CBA and C57Bl6 sperm is distinguishable to the trained eye, and makes a useful demonstration of the software’s features. The angle profiles generated are distinct for each genotype (Figure 2A). CBA sperm have a larger cross-sectional area, are longer, and also have slightly shorter hooks than C57Bl6 sperm (Figure 2B/C). These differences are reflected in the profiles; the long narrow tail in the CBAs appears as a smooth curve at x=50 in the profile, while the shorter, wider C57Bl6s show a distinct dip corresponding to the sharper curve of the dorsal angle before the acrosome. The shorter hook of the CBAs is also seen as a narrow peak at x=10; the longer hook of the C57Bl6s has a correspondingly wider peak. Automated segmentation of the nuclear profile allows quantification and significance testing of the inter-line differences in each separate region of the nuclear profile (see Supplementary figures 4, 5, 14).

CBA and C57Bl6 have previously been characterised by Wyrobek et al [9], who measured 160 nuclei of each genotype by manual tracing of projected microscope images of eosin-stained sperm heads. We found our measured values to be similar (Supplementary Table 7) but slightly smaller - as expected given that their measurements are for the entire sperm head rather than just the nucleus. The body widths are within 0.3μm, and our bounding heights are approximately 1.2μm smaller, consistent with our measurements lacking the acrosomal cap (~0.15μm [30]), and the proteinaceous part of the sperm hook. We measured the CBAs to be 12% longer than the C57Bl6s, again close to the previously published 13.5%.

**Figure 2:**
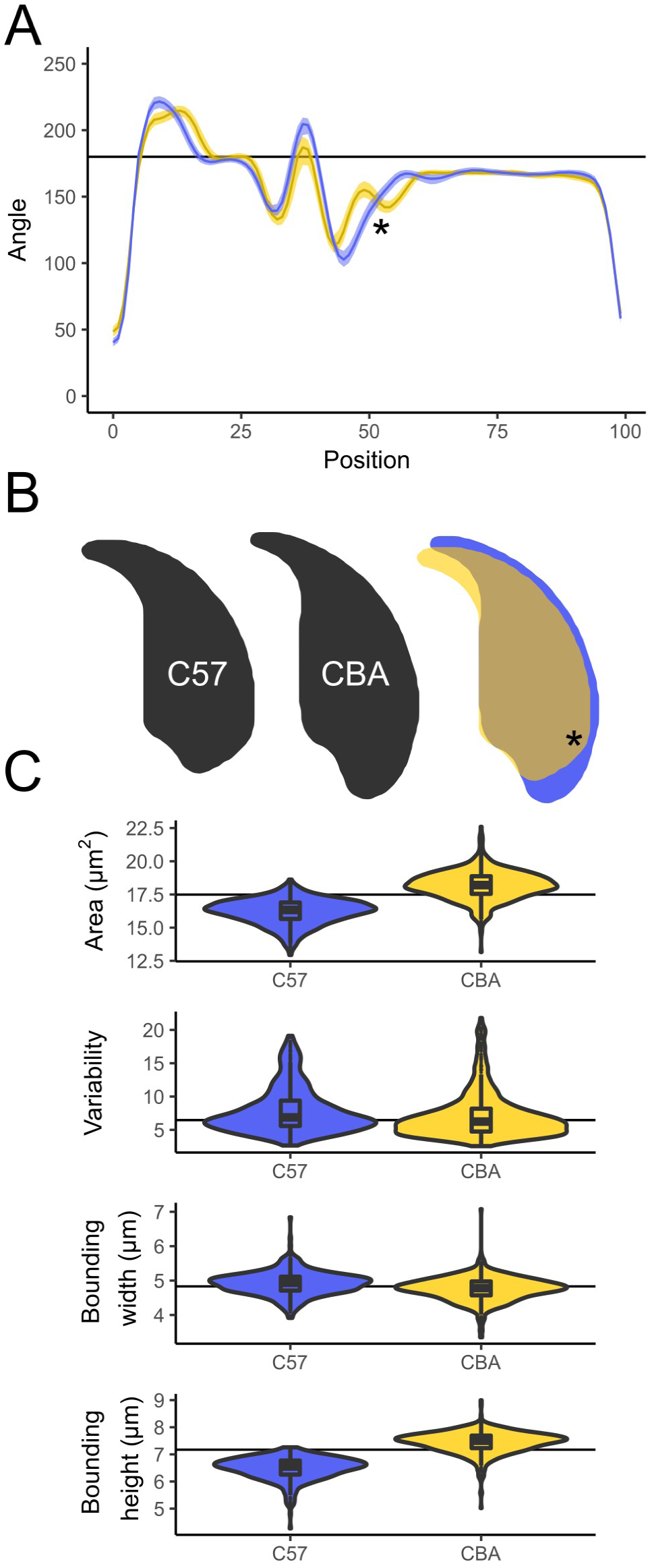
A) Comparison of shape profiles between C57Bl6 (yellow) and CBA (blue), showing the median and interquartile range of the nuclear shape profiles. B) Consensus nuclei from each population, and the overlap showing the regions differing. C) Size and shape measurements between the lines. The prominent dorsal angle in C57Bl6 nuclei is marked with an asterisk.

### Comparison of sperm morphology and variability across lines demonstrates the effects of inbreeding depression and hybrid dysgenesis

With the software tested on CBA and C57Bl6, we wanted to investigate the extent to which sperm shape variability within and between lines is affected by two factors: inbreeding depression and the complex inter-subspecific mosaic origin of classical laboratory strains. We selected a panel of inbred laboratory lines and compared them to (a) outbred laboratory lines, and (b) wild-derived inbred lines (Table 1). Biological replicate samples from the inbred lines represent either single animals (lab lines) or a pool of two animals (wild-derived inbred lines). For the outbred lines, several individuals were pooled to ensure we were capturing the diversity of the population as a whole.

Variability within each line was assessed using a new measure based on the similarity of each cell’s angle profile to the median for that line (see Methods). This was found to correlate well with other population measures of variability such as the coefficients of variation for area, bounding height and perimeter (Supplementary table 1). A comparison of the overlaid average nuclear shape is shown in Figure 3. In addition to each line having a characteristic sperm morphology, different lines showed different levels of intra-sample variability. A breakdown by biological replicates shows that these data reflect true line differences rather than biological differences between individual animals or technical differences between imaging sessions or choice of fixative (Supplementary figures 9-11).

**Figure 3:**
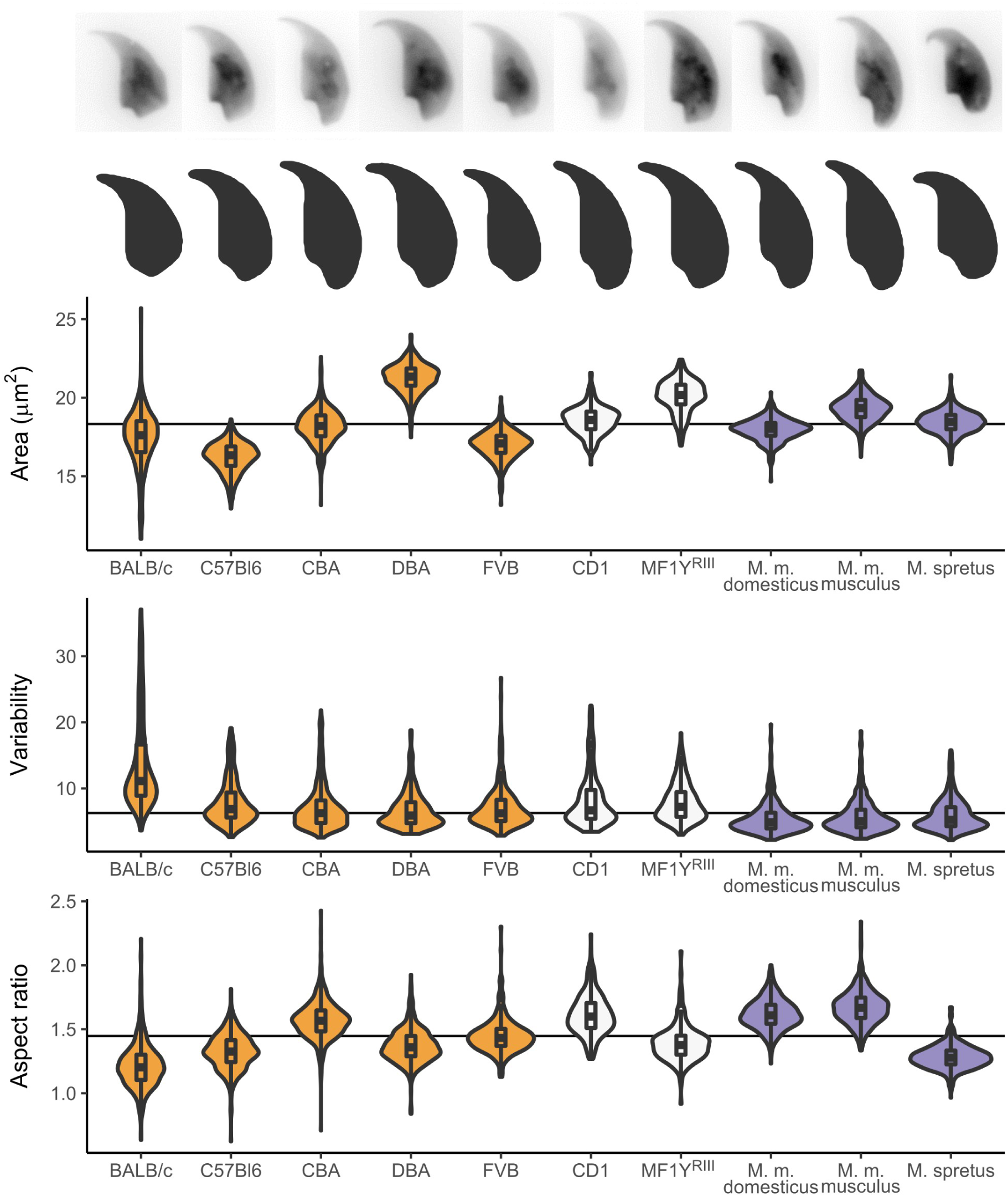
Parameters for additional lines examined, with representative nuclei and population consensus. Samples are coloured according to their type: from left to right: inbred (yellow), outbred (white) and inbred wild-derived (blue).

The BALB/c mice have the most variable shape profiles of all the lines we analysed, as well as the highest coefficient of variability in area, height and width (Supplementary tables 1, 2). The other inbred laboratory lines all showed low intra-line variability despite the fact that there were marked differences in sperm size and shape between lines. Of the inbred laboratory lines tested, CBA and DBA had the lowest intra-sample variability. The two outbred lines, CD1 and MF1Y^RIII^ both showed slightly higher intra-sample variability. This may reflect the fact that these samples were pooled samples derived from multiple genetically unique individuals. Turning to the wild-derived lines, all three lineages analysed (*M. m. domesticus, M. m. musculus* and *M. spretus*) had lower variability than any of the standard laboratory lines, despite that fact that these wild-derived lines are inbred.

### Segmentation of sperm profiles allows detailed analysis of elements of sperm morphology

Once “unrolled” into an angle profile, this profile can then be segmented at local minima and maxima (see Methods) to identify specific landmarks within the sperm head shape. Some landmarks are consistently found across all lines, such as the tip of the apical hook and the point of maximum curvature at the base of the sperm head, while other landmarks such as the dorsal angle and the indentation at the tail attachment site are variable between lines. As an example of how this can be used to compare samples, since we had already found the presence of a dorsal angle to vary between CBA and C57Bl6 sperm, we examined how this varied across the full data set. Of all the lines studied, only five showed a clear dorsal angle, with the others having a smoother profile posterior to the acrosome. The distance from the rear reference point to the dorsal angle was characteristic for each of these five lines, as was the variability in this measurement, with BALB/c mice showing highest variability. Supplementary Figure 15 discusses the ubiquitous and variable landmarks discovered by the segmentation analysis and shows the detailed segmentation pattern for each line, while Supplementary Table 4 gives the numerical segment length data for each line.

### C57Bl6 / CBA F1 cross males demonstrate the effects of each parental genotype on sperm shape and the relief of inbreeding depression by heterosis

The differences we saw between inbred and outbred laboratory lines made us curious as to the impact of line background and genetic interactions thereof. We investigated one specific reciprocal F1 cross, between C57Bl6 and CBAs. The use of F1 animals is important here as it relieves the effects of inbreeding depression caused by fixation of deleterious recessive variants in each of the parental lines, but still yields a uniform population of genetically identical males from each cross. B6CBA mice are the F1 offspring of a female B6 with a male CBA and CBAB6 mice are the reciprocal cross. Sperm morphology for both F1 lines matches the CBA parental line closely, indicating a dominant effect of the CBA genotype (Figure 4A). In terms of sperm cross-sectional area, both types of F1 sperm are much more similar to the CBA parent, while being fractionally larger than either parental line (Figure 4B). Males from both directions of the F1 cross showed less variability in their sperm shape compared to either parent line, suggestive of a degree of heterosis in the F1s.

**Figure 4:**
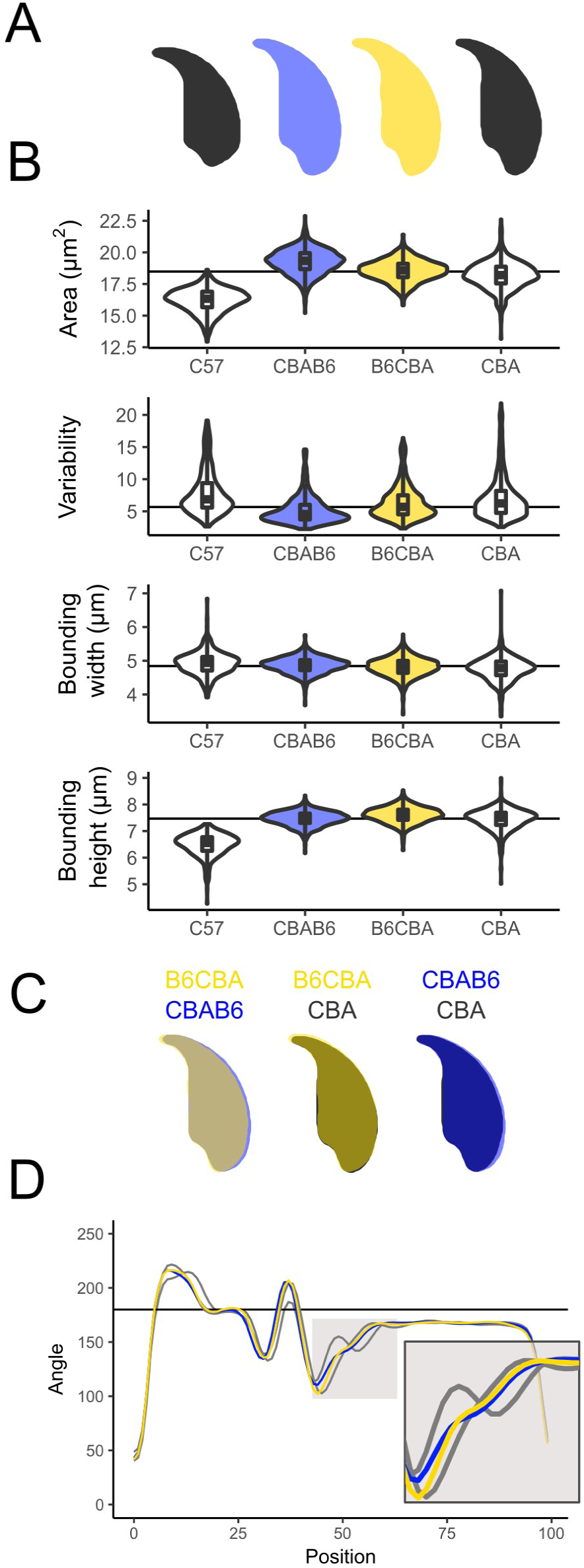
Subtle differences can be seen between a CBAB6 (CBA mother) and a B6CBA (C57Bl6 mother). Both are intermediate to the parental shapes, but CBAB6 sperm are wider, and their shape is closer to that of the C57Bl6. A) Consensus nuclei B) Size measurements; C) Overlay of consensus nuclei; D) comparison of angle profiles; the tail attachment region is expanded in the inset.

The reciprocal cross data allows us to look for parent-of-origin effects on sperm shape. We found two such differences, in sperm cross-sectional area and in bounding width. CBAB6s have a slightly larger sperm area than the B6CBAs (19.3 square microns versus 18.6 square microns, p<0.001) and the region around the posterior of the nucleus is widened in the CBAB6s, intermediate to CBA and C57Bl6 (Figure 4B/C). The differences around the posterior are largely driven by changes in the dorsal angle, which is present in C57, absent in CBA, and virtually absent in both reciprocal F1 cross males (Figure 4D). For bounding width, we find that this parameter is influenced by the male parent: CBAB6 and B6CBA are significantly different to each other (p=0.0016), as are C57Bl6 and CBA (p=1.27E-12), but there is no significant difference between C57Bl6 and CBAB6 (p=0.18) or between CBA and B6CBA (p=0.095). This suggests that this aspect of sperm shape may be influenced either by sex chromosome or mitochondrial background or by autosomal imprinted loci.

### Hierarchical clustering can separate samples based on shape differences

Next, we turned our attention to the analysis of morphological variation within a given population. In particular, we considered that cluster analysis of the sperm from a single sample would give an unbiased breakdown of the different morphological sub-populations contained therein. We used a hierarchical clusterer, as implemented by the WEKA data mining tool [29] to separate sperm based on their shape profiles.

We tested the clustering algorithm by pooling images from C57Bl6 and CBA and analysing them as a single sample. Since C57Bl6 and CBA sperm are slightly different sizes, the simplest partitioning of the mixed set is a binary cut-off at a given threshold for nuclear area. Passing the nuclear areas to a hierarchical clusterer and selecting the two most distinct clusters using the Ward clustering method was 83-85% accurate at separating the individual sperm by line. To determine whether shape-based hierarchical clustering could improve upon this, we sampled values from the angle profile for each nucleus at regular intervals (corresponding to the original window proportion) and provided these as inputs to the clustering algorithm. This clustering was markedly more accurate than a simple size-based cut-off, and separated the two genotypes with 91-95% success (Supplementary table 6). In head-to-head tests using a representative subset of 50 nuclei from each genotype, the clusterer performed at least as well (96%) as experienced assessors (97% accuracy), and substantially better than novice assessors (75% accuracy) (Supplementary figure 12).

### Hierarchical clustering can detect morphological subgroups within a sample

Having demonstrated that cluster analysis can recover different shapes from a mixed population of known composition, we looked at its use for novel shape discovery within a single highly variable population. Since the BALB/c line showed the highest variability in our line survey, we chose this as our test sample. A cluster analysis based on angle profile alone found four major groups of sperm shape, from mostly normal through to severe hypercondensation of the sperm (Figure 5). The final class is still highly variable compared to the other classes; clustering these nuclei further reveals a separation of two separate types of hypercondensation (Supplementary figure 16) as previously described [31]. While the most normal sperm had near-normal placement of the dorsal angle, and a normal tail attachment site, the most heavily distorted sperm showed frequent presence of additional sharp angles in the sperm outline, effacement of the tail attachment site due to compression of the rear of the sperm head, and an ever more prominent and misplaced dorsal angle that may reflect altered microtubule dynamics during nuclear shaping (see Discussion).

**Figure 5:**
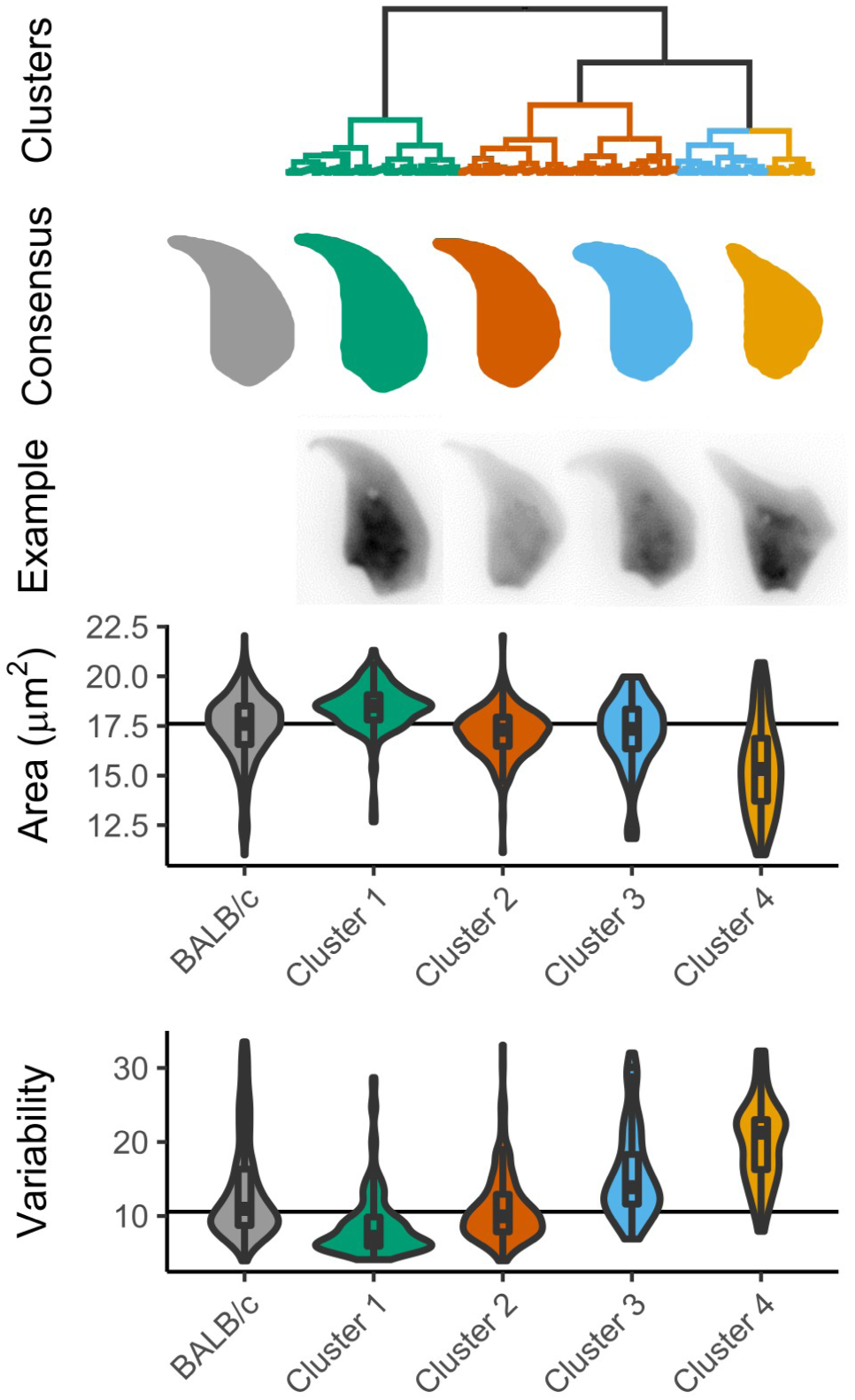
The overall population of BALB/c sperm appears distorted compared to other lines (grey), but clustering reveals separate classes of morphology, from mostly normal (green) to highly condensed (yellow).

## Discussion

We present here a morphological analysis tool designed to study nuclear morphology, with the ability to automatically identify key landmarks in the nuclear outline and quantitatively measure a range of nuclear and sub-nuclear parameters. Here, we demonstrate the use of this software to analyse the highly asymmetrical shape of the mouse sperm nucleus; however it is a generally applicable tool suitable for analysis of all sizes and shapes of nuclei. A companion paper ([32], submitted for publication) demonstrates its use in comparing sperm from boars judged to be suitable/unsuitable for use in artificial insemination.

### Comparison of this method with other nuclear shape analysis methods

The key advantage offered by the software presented here is automation of the steps involved in object detection, shape decomposition and comparison. At the object detection stage, we use an edge detection algorithm that is markedly more effective than the fixed-threshold detection used in other packages, particularly in the presence of inhomogeneous staining of the bright chromocenter and dim apical hook. At the shape decomposition step, we introduce a modification of the Zahn-Roskies transform [28] that sensitively detects the various angular landmarks around the sperm periphery without the need for manual intervention. Together, these innovations massively increase the number of nuclei that can be quantified and compared to each other, with a total of 8,749 nuclei being measured during this study, and over 22,000 nuclei in the companion paper analysing boar sperm [32]. This for the first time permits the use of sample sizes that accurately capture not only fixed size and shape differences between samples, but also the detection and classification of intra-sample variability. Our method is robust to differences between camera and microscope setups and fixation techniques, making it amenable to analysis of large numbers of images, and potentially to automated image capturing from whole slide scanners.

While there are other features of sperm morphology that we do not yet address in this package, the modular design of our software allows additional analysis pipelines to be added at a subsequent date, and for features from different fluorescence channels to be associated with specific nuclei and analysed in relation to them. We anticipate that other sperm morphological features such as the extent and thickness of the acrosome, the proteinaceous tip of the hook, the presence of cytoplasmic droplets, and the length and morphology of the tail will be amenable to our approach by combining nuclear staining for orientation with phase contrast imaging, tubulin immunostaining, MitoTracker, SpermBlue or other stains. Since we are imaging fixed cells, the nuclei also remain available for interrogation by chromosome painting or other molecular cytogenetic approaches, e.g. to detect aneuploid cells and correlate their chromosomal status with their nuclear morphology.

### Comparison of sperm shape within and between lines

Our observations support previous studies (e.g. [4,9]), add further information on the precise regions of the sperm head that that differ between lines, and demonstrate the variability of sperm morphology within each given line. In particular, we examined the presence and placement of the dorsal angle of the sperm. This feature is created by pressure from the manchette: a cone-shaped array of microtubules that forms behind the nucleus and slides backwards during spermiogenesis, shaping the rear of the sperm head in the process. Defects in katanin p80, a microtubule severing protein, lead to failure of this process and abnormal compression of the base of the sperm head [6]. The narrowing of the tail attachment site seen in FVB and BALB/c males, together with the prominent dorsal angle seen in both lines (especially the latter) may indicate that manchette migration is abnormal in these males.

### Comparison of sperm variability within and between lines

The greatest variability we saw was in the BALB/c animals. This line is known to have poor sperm morphology and high levels of sperm aneuploidy. Kishikawa et al [31] observed different classes of abnormality, which we were able to recapitulate. In their analysis, the authors found chromosomal abnormalities in 35% of highly abnormal sperm, but also in 15% of sperm that were morphologically ‘normal’ by their criteria. Given that our new analysis detects additional classes of more subtle shape difference that were not discriminated in the earlier analysis, we hypothesise that these new abnormal classes may also be enriched for chromosomal defects compared to the most normal sperm. Further differences await characterisation: different classes and levels of sperm abnormalities have been described depending on the particular subline and age of the animal [33].

Consistent with [34], we found that an F1 cross between C57Bl6 and CBA laboratory lines lowered sperm shape variability (see below), suggestive of a degree of inbreeding depression that was relieved by heterosis. However, the least variable lines we examined were the wild-derived inbred lines PWK, LEW and STF, representing *M. m. musculus, M.m. domesticus* and *M. spretus* respectively. Since these three lines are also inbred, this suggests that the wide variety of sperm shapes in laboratory lines, and the elevated level of intra-individual variability in all the laboratory lines is not primarily a consequence of inbreeding depression.

Instead, this is potentially linked to the status of the laboratory mouse as a hybrid between several mouse subspecies - a factor that may have disrupted regulatory interactions throughout the genome. Against this, PWK, despite being predominantly of *musculus* origin, nevertheless has substantial introgression of domesticus DNA, of the order of ~6-7% of the genome [35,36]. The degree of disruption may therefore depend on both the direction of introgression and the specific regions involved. The recent finding of polymorphic hybrid incompatibilities within both *musculus* and *domesticus* subspecies shows that multiple regions of the genome contribute to hybrid breakdown and hybrid sterility. Consequently, the various different classical and wild-derived inbred lines may have fixed different combinations of incompatible alleles that collectively destabilise sperm development to varying extents in each line [37].

X/Y mismatch is a strong potential contributor to regulatory disruption, since most laboratory lines carry a *musculus* Y on a predominantly *domesticus* background [35]. The copy number of the ampliconic genes on the X and Y chromosomes varies markedly between *musculus* and *domesticus* subspecies, and the relative copy number of these genes is known to be important for normal sperm morphology [38–40]. However, while most of the laboratory lines we examined do indeed have mismatched X/Y chromosomes [35,41,42], the FVB X and Y are both of *domesticus* origin, indicating that the alterations in sperm shape in this line are not due to X/Y mismatch.

An alternative but not mutually exclusive explanation for the difference between classical laboratory inbred lines and wild-derived inbred lines is that the classical lines have been selected over multiple generations for their ability to breed well in captivity - indeed FVB is particularly known for its fecundity [43]. It may seem paradoxical that selection for high fecundity could adversely affect male fertility parameters: however, under laboratory conditions of non-competitive mating, co-housing a single male with one or more females, it is likely that reproductive output is driven largely by maternal factors. Thus, even though laboratory lines are fertile under lab breeding conditions, their sperm may be uncompetitive in mixed mating experiments compared to a pure species background. The morphology of the FVB zygote pronucleus is independent of the paternal genetic background, and the efficiency of FVB sperm for IVF appears unexceptional [44]. Sperm morphology and fertilisation success in laboratory mice has been shown to evolve rapidly in response to competitive mating experiments, indicating that the baseline competitive ability of laboratory line sperm is sub-optimal [45,46]. Intriguingly, it has even been shown that in lines experimentally selected for high fecundity, male fertility and sperm morphology/motility parameters are compromised, suggestive of a trade-off between the male and female factors necessary for high fecundity in a laboratory environment [47].

### Relevance for speciation, fertility, and toxicology studies

Abnormal sperm head morphology has emerged as a common form of hybrid male sterility in mice [21–24,48]. Some sterility factors broadly impair spermatogenesis, resulting in reduced sperm counts, lower motility, and abnormal morphology. However, several studies have now shown that hybrid sterility QTL in mice often correspond to specific reproductive phenotypes [24]. The challenges of manually quantifying morphology in large mapping panels has necessitated the use of crude categorical scores [21,23,48], hampering quantitative precision and likely limiting the ability to draw causal links between hybrid incompatibilities and specific aspects of sperm morphological development.

Our approach assists in two ways: firstly by enabling more rigorous quantitation of sperm shape, and secondly by enabling the large sample sizes and systematic approach needed for mapping studies. As a proof of principle, we have compared males from a reciprocal cross between C57Bl6 and CBA mice, and identified a dominant effect of the CBA genotype on sperm shape. Within this, however, there are subtle differences between the CBAB6 and B6CBA animals, suggesting an effect of either chromosome constitution or imprinting on sperm bounding width. This demonstrates the usefulness of this approach for understanding subtle features of mouse sperm nuclear development, and the potential to use this software for genetic mapping of the various determinants of mouse sperm head shape.

Fertility rate and IVF efficiency has been correlated with the genetic background of sperm among inbred mouse lines [49]. Furthermore, many studies have shown that the genetic background of a line can influence sperm morphology. For example, deletion of the long arm of the Y chromosome results in a more severe phenotype on B10.BR background than on CBA [50]. Mashiko et al [16] have suggested morphology of sperm is associated with fertilising efficiency in at least two mouse lines (B6D2F1 and C57Bl6/N). Since particular genetic mutations in mouse sperm shape are associated with characteristic nuclear shape abnormalities (e.g. [19]), detailed examination of sperm from natural mutant and/or targeted knowckout animals may point to pathways of interest for understanding spermiogenesis and male fertility more generally.

In toxicological analysis, rodent sperm are conventionally manually classified into classes of predefined morphological abnormality (e.g. [26,51]). The hierarchical clustering implemented within the software is able to separate nuclei based on shape as accurately as an experienced manual sperm scorer; however it is much faster and more consistent. This may be of use in samples where the nature and degree of abnormalities is hard for humans to reliably quantify. It is also important to understand and quantify normal morphological variation between lines since different lines can have different responses to toxicological agents [52]. While many studies of toxicology using rodent models are conducted on rats, the extra information available in the mouse sperm head still makes them a useful model system. The fact that specific genetic lesions cause specific shape changes means that the sperm shape might in principle give information not just about the presence/absence of toxicity but also its mode of action. This level of analysis would complement existing studies of sperm function, which, in clinical settings or in automated CASA platforms (e.g. [53]), is still lacking detailed morphological data [10].

## Conclusions

We present a new software package for the rapid, high-throughput, replicable analysis and comparison of nucleus shape in mouse sperm. By using a range of mouse lines, we have demonstrated the ability of the software to discriminate subtle differences between lines, and to reproducibly separate the nuclei into morphological groups. This has applications for studies of speciation, fertility and understanding the impact of genotoxic compounds. The analysis steps are generalisable and will work on many symmetric or asymmetric shapes of nuclei including, but not limited to sperm from other species.

## Acknowledgements

We thank the animal handling staff at the University of Kent, University of Cambridge, University of Montana and Charles River Laboratories. We also thank the experimental test subjects who volunteered to classify C57Bl6 and CBA sperm.

## Authors’ contributions

Conceptualisation, BMS and PE; Methodology, BMS and PE; Software and Validation, BMS; Investigation, CCR, JB, EEPJ and GY; Data Curation and Formal Analysis, BMS and CCR; Visualisation, BMS; Supervision and Project Administration, PE; Writing - Original Draft, BMS and PE; Writing - Review and Editing, BMS, CCR, JMG, ELL and PE; Resources, JMG, ELL, EEKK, NA and PE; Funding Acquisition, NA and PE. All authors gave final approval for publication.

